# Contributions of Implicit and Explicit Memories to Sensorimotor Adaptation of Movement Extent During Goal-Directed Reaching

**DOI:** 10.1101/2020.10.22.350645

**Authors:** Devon D Lantagne, Leigh Ann Mrotek, Rebecca A Slick, Scott A Beardsley, Danny G Thomas, Robert A Scheidt

## Abstract

We examined how explicit memory of kinematic performance contributes to motor adaptation during goal-directed reaching. Twenty subjects grasped the handle of a horizontal planar robot that rendered spring-like resistance to movement. Subjects made rapid “out-and-back” reaches to capture a remembered visual target at the point of maximal reach extent. The robot’s resistance changed unpredictably between reaches, inducing target capture errors that subjects attempted to correct from one trial to the next. The subjects performed four sets of trials. Three were performed without concurrent visual cursor feedback of hand motion. Two of these required self-assessment of performance between trials, whereby subjects reported peak reach extent on the most recent trial. This was done by either moving a cursor on a horizontal display (visual self-assessment), or by moving the robot’s handle back to the recalled location (proprioceptive self-assessment). Two control conditions were performed either without or with concurrent visual cursor feedback of hand motion. We analyzed movement kinematics and used regression analyses to quantify the extent to which prior reach errors and explicit memories of prior performance contribute to subsequent reach performance. Consistent with prior reports, providing concurrent visual feedback of hand motion increased reach accuracy and reduced the impact of past performance errors on future performance, relative to the no-vision control condition. By contrast, we found no impact of interposed self-assessment on reach kinematics or on how prior target capture errors influence subsequent reach performance. Self-assessments were biased toward the remembered target location, differing markedly from actual reach performances. Including self-assessments as explicit memory of performance in the regression analyses did not improve model predictive accuracy. Therefore, we conclude that explicit memories of prior kinematic performance do not contribute to implicit sensorimotor adaptation of movement extent.

## Introduction

The human sensorimotor system is adept at performing goal-directed actions in the presence of changing environmental conditions due to the brain’s remarkable ability to compensate for performance errors that arise during movement (Shadmehr and Mussa-Ivaldi 1994; Lackner and Dizio 1994; Shadmehr and Brashers-Krug 1997). Even in the simplest actions such as reaching, corrections for performance errors are comprised of separate components attributable to implicit sensorimotor adaptation (Shadmehr and Mussa-Ivaldi 1994; Thoroughman and Shadmehr 2000; Scheidt et al. 2001; Izawa et al. 2008; Judkins and Scheidt 2014) and explicit strategic re-aiming (Mazzoni and Krakauer 2006; Taylor and Ivry 2011; Taylor et al. 2014) (see also Redding and Wallace 1996). Both implicit and explicit mechanisms utilize memories of prior performance features to improve subsequent performance. Implicit learning occurs subconsciously such that individuals may be unaware that they have altered their behavior and/or cannot consciously verbalize strategies used to adapt (Frensch 1998; Mazzoni and Krakauer 2006). Implicit learning can occur automatically without explicit knowledge of results (Magescas and Prablanc 2006; Prablanc et al. 2007). By contrast, explicit learning refers to strategic changes in behavior guided by conscious decisions (Magill 2011; Taylor et al. 2014; Heuer and Hegele 2015; Krakauer et al. 2019). This type of learning can be assessed by asking people to report information about upcoming plans for movement or about features of a completed movement.

Prior studies have sought to understand how explicit and implicit processes might interact during motor learning. Boyd and Winstein (2003) investigated the effect of explicit information on implicit learning in individuals with focal stroke affecting regions of the brain that mediate information transfer between explicit and implicit memory. They found that using explicit information to improve performance in a serial reaction time task had a negative impact on implicit learning, suggesting that the explicit information may have hindered the use of the implicit memories. Similar results were seen in healthy individuals during a visual motor rotation task by Benson et al. (2011), who suggested that the results reflect a competition between explicit and implicit learning mechanisms for a limited spatial working memory resource. By contrast, others have reported that implicit learning can interfere with the effectiveness of explicit error correction. Mazzoni and Krakauer (2006) set up a conflict between implicit and explicit error correction by instructing subjects to the exact nature of directional perturbations in a visual motor rotation learning task and how they could compensate for it. Although subjects initially succeeded in compensating the imposed rotation with the suggested strategy, they made increasingly large errors with practice. The investigators concluded that the explicit strategy of re-aiming was undermined by implicit learning of the rotation. They also found that the rate of implicit learning was the same for subjects informed of the explicit strategy and for those who were not, indicating that explicit planning did not interfere with implicit learning in that task. Relatedly, Reber and Squire (1994, 1998) showed that patients with deficits in episodic memory performed well on a serial reaction time task, implying that this component of explicit memory is distinct from that required for implicit learning. Altogether, these studies suggest that while implicit and explicit processes can both contribute beneficially to motor performance, their contributions are likely distinct and interfere with one another in complex, task-dependent ways.

Here, we investigated how explicit and implicit memories contribute to reach adaptation. Subjects grasped the handle of a planar robot while making rapid out-and-back reaches to a single visuospatial target. The robot opposed motion with spring-like loads that changed from one reach to the next. Between movements, we sometimes required subjects to recall and report the hand’s location at the moment of peak movement extent on the most recent trial; these self-assessments were used to provide visibility into explicit memory of kinematic performance. We used multilinear regression to fit models of sensorimotor adaptation to data from trial blocks performed with and without self-assessments to compare the contributions of implicit and explicit memories to the trial-by-trial compensation for variable environmental loads. We evaluated whether interposed self-assessments influence the way people use sensorimotor memories during adaptation and we tested the hypothesis that implicit and explicit memories both contribute meaningfully to the adaptation of kinematic performance during repeated practice of goal-directed reaching. Preliminary aspects of this work have appeared in abstract form (Slick et al. 2017).

## Methods

### Participants

Twenty healthy subjects [mean age 25.5 ± 6.1 years (mean ± SD here and elsewhere); 13 females and 7 males] provided written, informed consent to participate in this study. All had normal or corrected-to-normal vision, and none had any known neurological deficits. Subjects were recruited from the Marquette University campus community. All experimental procedures received institutional approval in accordance with the Declaration of Helsinki.

### Experimental Setup

Each subject sat in a high-backed chair and grasped the handle of a horizontal planar robot with the right hand (Fig 1A). An overhead sling supported the right arm against gravity. The robot was actuated by two brushless DC torque motors (M-605-A Goldline; Kollmorgen, Inc. Northampton, MA). Handle location was resolved within 0.038 mm using joint angular position data from two 17-bit, encoders (A25SB17P180C06E1CN; Gurley Instruments, Troy, NY). Robot control and data collection were performed at 1000 samples per second. Handle kinematic data and robot control signals were stored to disk for post-processing.

**Figure 1:**
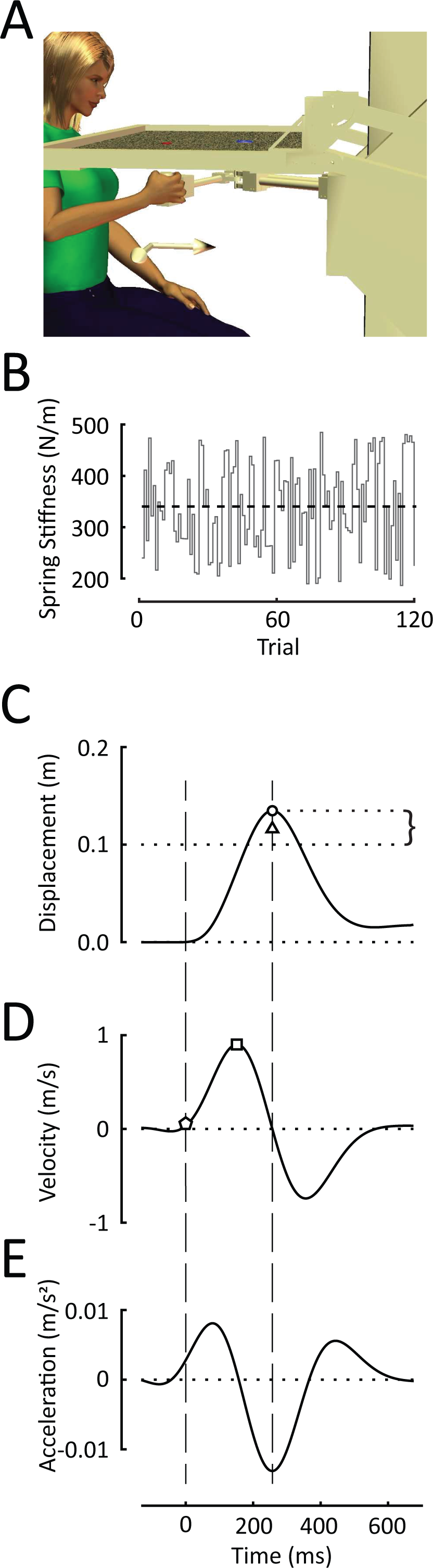
A) Experimental setup: Home and goal targets were projected on a horizontal screen that blocked the subject’s view of the hand, arm and robotic manipulandum. B) The magnitude of the robot’s spring-like load varied pseudorandomly from trial to trial. Dashed line: mean spring stiffness across all trials. Panels C, D, and E) Hand displacement, velocity, and acceleration for a typical reach trial. Circle: maximum movement extent; Brace: reach error; Triangle: self-assessed maximum movement extent; Square: peak velocity; Pentagon: 10% of peak velocity (i.e., movement onset); Dashed vertical lines mark the movement onset and target capture times.

A horizontal display screen was mounted 2 cm above the robot handle to occlude vision of the arm and hand as subjects performed goal-directed reaching movements. A starting “home” target was projected onto the display screen ∼28 cm anterior of the subject’s right shoulder. A goal target was placed 10 cm further from the home target in the sagittal plane. A scintillating random dot field (60 cm high x 30 cm wide; 30 Hz refresh rate) was projected on the screen at all times to minimize the impact of extraneous visual cues on performance of the tasks described below. Unless indicated otherwise, no ongoing visual feedback of hand motion was provided during the tasks.

During reaching, the robot rendered a spring-like load that resisted hand motions away from the home target. No force was applied when the handle was centered on the home target. The strength of the spring-like load changed unpredictably from one movement to the next (i.e., from trial to trial), but did not change within a trial (Fig 1B). The sequence of loads was drawn from a uniform distribution that was constructed to have negligible autocorrelation structure. The load sequence had a mean stiffness of 339 N/m and a standard deviation of 89 N/m. All subjects experienced the same sequence of perturbations.

The motors also rendered a stiff mechanical channel that constrained hand motion to the sagittal plane (c.f., Scheidt et al. 2000), effectively limiting performance variability to that of movement extent.

### Procedures

Subjects performed four blocks of 120 goal-directed movement trials wherein they were instructed to “Reach out-and-back in one fluid motion to hit the remembered goal target at the peak of your reach.” A trial began when a “GO!” cue was projected onto the visual display screen; both the home and goal targets disappeared at this time. Subjects were to perform a ballistic, 10 cm out-and-back movement to hit the remembered goal target. After the reach was completed, the subject relaxed as the robot slowly and smoothly re-centered the hand at the home position. During the re-centering, the hand’s cursor and the home and goal targets were visible to minimize the impact of “proprioceptive drift” on reach planning (cf., Wann and Ibrahim 1992). After a variable relaxation interval (750 ms to 825 ms) a new trial began. In each of the four blocks, the first 20 reaches were considered practice trials, which allowed the subjects to become familiar with the task and the desired movement extent. During practice trials, concurrent visual feedback of hand position (i.e., the hand cursor) was projected onto the horizontal screen during the reach. Immediately following the practice trials, subjects performed 100 test trials, which we used to characterize how memories of prior reach performances influenced subsequent reach performance under four different experimental conditions.

Each block contained a different condition depending on whether visual information was provided about ongoing hand motion and whether or not the participant was asked to explicitly report the outcome of the most recent performance. In the no visual feedback, no assessment (NV-NA) control block, subjects performed the 100 test reaches without concurrent visual cursor feedback and without interposed self-assessments. In the no vision, proprioceptive assessment test block (NV-PA), subjects performed the 100 test reaches without concurrent cursor feedback, but with proprioceptive self-assessment interposed between each reach, as described below. In the no vision, visual assessment test block (NV-VA), subjects performed the 100 test reaches without concurrent visual feedback but with visual assessment between reaches. In the visual feedback, no assessment (V-NA) contrast condition, subjects performed the 100 test reaches with concurrent visual feedback and without interposed self-assessments. Providing concurrent visual feedback of endpoint motion is known to impact how sensorimotor memories contribute to subsequent reach attempts (Judkins and Scheidt 2014). We included this last block to provide a contrast condition with which to compare the results of the self-assessment conditions. Each block took about 20 minutes to complete. Block order was counterbalanced across subjects to reduce potential order effects.

In the two blocks that included interleaved self-assessments, subjects were to explicitly report the maximum extent of hand movement after each reach. These self-assessments provide an objective measure of explicit memory of task performance. In the self-assessment blocks, subjects first reached out- and back to the remembered goal target without concurrent visual feedback of hand or cursor motion (Reaching). They then pointed to the location of peak movement extent on the previous reach in one of two ways (Pointing; i.e., post-reach performance reporting). One test block required “proprioceptive assessment” (PA), whereas the other used “visual assessment” (VA). For proprioceptive self-assessments, subjects were to recall and point to the remembered location of maximum reach extent by actively moving the robot handle to that remembered location. The robot’s motors were disabled during self-assessments such that subjects could not use hand forces to infer hand displacement. No cursor feedback or visual landmarks were provided. The scintillating random dot field discouraged visual fixation about the remembered target location as the subject moved his or her hand to report the point of maximum reach extent. Subjects pressed a “Select” button with their left hand to confirm that location. For visual self-assessments, subjects pointed to the location of maximum reach extent by moving a cursor projected onto the screen with a random initial location along the line between initial and goal targets. Subjects repositioned the cursor by pressing “Closer” and “Farther” buttons with their left hand. To confirm the location of maximum reach extent, subjects pressed a third “Select” button with their left hand. Only the cursor and scintillating field were visible during visual self-assessments. After each self-assessment, the subject relaxed as the robot re-positioned the hand back to the home position in anticipation of the next reach trial.

### Data Analysis

All trials were visually inspected prior to post-processing. Trials were precluded from analysis for one or more of the following reasons: if the hand drifted 1 cm or more from the home position before the “GO!” cue; if the movement was not ballistic (i.e., if the time between movement onset and peak movement extent exceeded 500 ms); or if the reach was not directed along the channel constraints, causing robot motor torques to exceed predefined safety limits. Across subjects, only 2.8% ± 5.1% of evaluation trials were excluded from further analysis.

We quantified *reach error* (*ε*_*i*_) as the signed difference between the maximum extent of the out-and-back-reach and the actual target distance of 10 cm (Fig 1C, brace). Positive values indicated overshoot of the target. We quantified *reach precision* (*σ*_*ε*_) as the standard deviation of the reach errors across evaluation trials within each block. We defined *movement onset* for each reach as the first moment where hand speed exceeded 10% of its first peak value (Fig 1D, pentagon). We quantified *movement time* as the difference between movement onset and the time of the maximum reach extent (c.f., time between vertical dashed lines in Figs 1C-E). We quantified *assessment error* 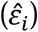 for the proprioceptive and visual assessment trials as the difference between the self-assessed peak movement extent (Fig 1C, triangle) and the actual target distance.

### Models of Sensorimotor Adaptation

We next investigated how implicit and explicit memories of reach performance contribute to the adaptive response to changing environmental loads. We used step-wise linear regression to fit a family of linear, “fast process” adaptation models (Smith et al. 2006; Lee and Schweighofer 2009; Judkins and Scheidt 2014) to the time series of reach errors (*ε*_*i*_,) within each block of trials. These models considered four independent input variables. We regard the observed movement extent on the previous trial (i.e., *ε*_*i-*1_) as a proxy for implicit memory of reach performance (c.f., Thoroughman and Shadmehr 2000; Scheidt et al. 2001; Judkins and Scheidt 2014). We regarded self-assessed movement extent 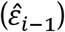 as a proxy for an explicit (declarative) memory of reach performance. We also account for the impact of the robot’s physical resistance on movement by including input terms reflecting the current trial’s spring-like load (*k*_*i*_), as well as a memory of the robot’s most recent load (*k*_*i-*1_). All variables were centered such that their respective means were subtracted from the time series prior to model fitting. The criteria for a term to enter the model was p ≤ 0.05. The criteria for a term to be removed from the model was p ≥ 0.10.

Multilinear regression analyses can be sensitive to multicollinearity within the set of input variables. Even though we designed the sequence of loads *k*_*i*_, to be minimally correlated with the sequence of prior trial loads *k*_*i-*1_, we expect structural multicollinearity to arise from mechanical interactions between the robotic spring-like loads and the inherent compliance of the subject’s arm. Subjects are expected to make shorter reaches when the robot renders a stiffer spring than when it renders a more compliant spring. Consequently, we expect the sequences of kinematic memories (i.e., *ε*_*i-*1_ and 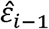) to correlate strongly with the sequence of prior spring-like loads *k*_*i*_. To address this known source of collinearity, we derived a new set of input variables for use in our stepwise regression analyses. These include the original load sequences *k*_*i*_ and *k*_*i*-1_ as well as two new “residual” sequences *e*_*i*-1_ and *ê*_*i-*1_ that were derived from the original memory sequences *ε*_*i*_ and 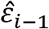 by subtracting out all linear dependence on the sequence of prior loads *k*_*i*-1_. By managing multicollinearity in this way, the stepwise regression analyses can yield insight into how much variability in reach performance depends on the robotic perturbations, and how much depends on the unique information contained within the sequences of implicit and explicit memory.

As a basis for comparison, we also considered a model of sensorimotor adaptation examined in several prior studies of horizontal planar reaching (Scheidt et al. 2001; Judkins and Scheidt 2014):

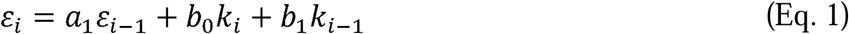

Here, the model coefficients (*a*_1_, *b*_0_, and *b*_1_) describe how reach performance is influenced by a memory of prior reach errors, and by the current and previous loads, respectively.

### Statistical Hypothesis Testing

This study addressed three questions. First, we asked whether engagement of the explicit memory system during the recall and reporting of recent reach performance would influence sensorimotor adaptation on subsequent reach attempts. To this end, we used one-way repeated measures ANOVA and Dunnett post-hoc *t*-test to compare (relative to the NV-NA control condition) the extent to which interposed self-assessments impact movement kinematics. Second, we asked whether memories of prior performances recalled during self-assessment accurately reflect peak movement extents performed on prior reaches, or whether explicit memories of performance may differ systematically from objective performance. To do so, we used linear regression analyses and planned *t*-tests to evaluate slope and bias in the relationship between self-assessed and actual movement extents for both the proprioceptive and visual self-assessment conditions. Third, we wished to determine the extent to which information encoded by the explicit memory system contributes to sensorimotor adaptation in able-bodied individuals. We used stepwise regression to fit the {*ε,k*,*e*,*ê*} datasets in the NV-PA and NV-VA blocks to determine whether the additional information about explicit memories ^ provided by visual and/or proprioceptive self-assessment would contribute meaningfully to kinematic performance on subsequent reaches. All data processing and model fitting were done in MATLAB 2019a (The MathWorks, Natick, Massachusetts). All statistical analyses were conducted using the SPSS statistics software package (IBM corp. Armonk, New York). Statistical significance was set at a family-wise error rate of *α* = 0.05.

## Results

All subjects were attentive throughout the experimental sessions and completed all four blocks of trials. Hand trajectories had similar kinematics across all four blocks, as shown in Figure 2 for a selected subject. As instructed, the subject reached briskly out-and-back to the approximate location of the remembered target. The subject did not pause at the time of peak movement extent, which averaged 239 ± 14 ms across all conditions and did not exceed 322 ms from the moment of movement onset in any trial. Reach error for this subject averaged 2.46 ± 1.47 cm across all conditions. Velocity profiles were bi-phasic and acceleration profiles were consistent with ballistic reaching (i.e., without evidence of mid-reach corrections). Performance varied trial-by-trial because the robot rendered mechanical loads that varied trial-by-trial. Consequently, movement kinematics substantially overlapped across all four experimental conditions. The subject’s performance shown in Figure 2 was typical of the study cohort. Movement times averaged 256 ms across conditions, with an average standard deviation of 21 ms within the study cohort. Reach errors averaged 2.49 cm across conditions, with an average standard deviation of 1.92 cm within the cohort. Thus, movement patterns were consistent within subjects across all testing conditions.

**Figure 2:**
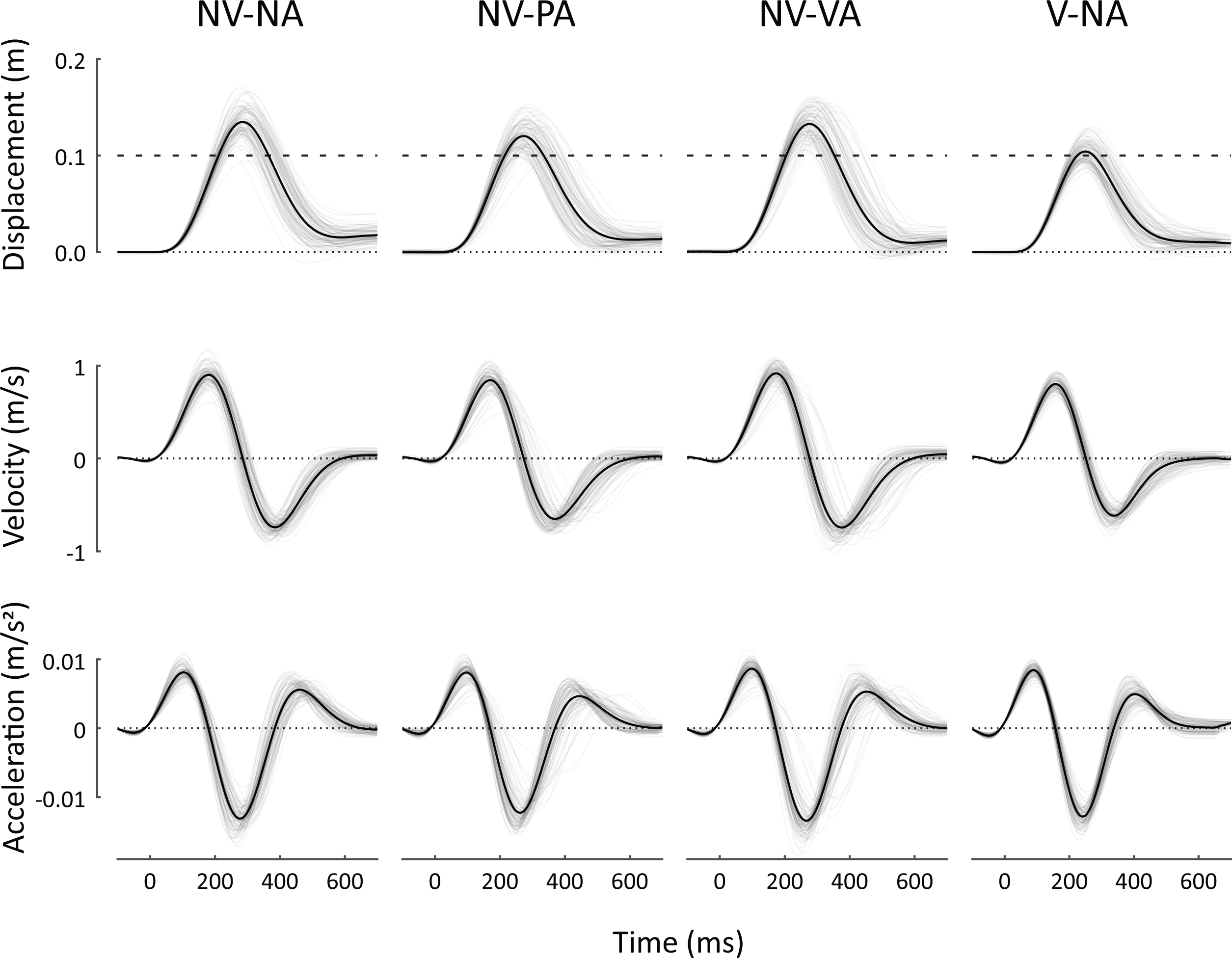
Overlaid reach trajectories in all conditions for a selected subject; each column corresponds to a single test condition. NV-NA: no vision, no assessment control condition; NV-PA: no vision, proprioceptive assessment; NV-VA: no vision, visual assessment; V-NA: concurrent vision, no assessment contrast condition. Top: displacement vs. time. The dashed horizontal line at 10 cm represents the target location. Middle: hand velocity. Bottom: hand acceleration. Thin gray lines: individual trials. Bolded lines: the average of all trials.

Within the cohort, movement timing overlapped to a large extent across the four experimental blocks. The across-subject average *range* of durations *within* each of the four conditions was greater than 75 ± 28 ms (V-NA) but less than 106 ± 39 ms (NV-PA). By contrast, the difference in average movement durations averaged 45 ± 26 ms *across* conditions, a fraction of the range of movement times within each block of trials. We observed small but systematic differences in the extent of hand position at target capture dependent on the presence or absence of concurrent feedback of cursor motion (Fig 2A; V-NA compared to other conditions). The across-subject average *range* of movement extents within each of the four conditions was greater than 5.2 ± 0.8 cm (V-NA) but less than 8.1 ± 3.4 cm (NV-PA). Large ranges are to be expected because the robot changes its resistance to movement from one trial to the next. By contrast, the *across* condition difference in average movement extent did not exceed 3.4 cm for any subject, again a fraction of the range within each block of trials. Thus, movement kinematics were consistent in the sense that movement durations and extents both overlapped substantially across the four conditions.

### Impact of vision and self-assessment on sensorimotor adaptation - movement kinematics

We observed no interference between the reach and self-assessment tasks with regards to the kinematics of reaching. One-way repeated measures ANOVA found a main effect of testing condition on the mean reach error at peak movement extent [F_(3,57)_ = 19.04, p < 0.0005; Fig 3A] and on the variability of reach extent [F_(3,57)_ = 10.50, p < 0.0005; Fig 3B]. These effects were due predominantly to the presence of visual feedback of cursor motion during reaching; Dunnett’s post hoc *t*-tests confirmed that movements in the V-NA contrast condition were more accurate (T_19_ = 5.51; p < 0.0005) and less variable (T_19_ = 3.73; p = 0.001) than those in the NV-NA control condition. By contrast, movement accuracy and precision in the two self-assessment blocks did not differ systematically from those in the NV-NA control condition (T_19_ < 1.42 and p > 0.05 in all cases). Thus, providing a visual representation of the moving hand improved reach performance relative to the NV-NA control condition, but interposing self-assessments between reaches neither improved nor degraded reach performance.

**Figure 3:**
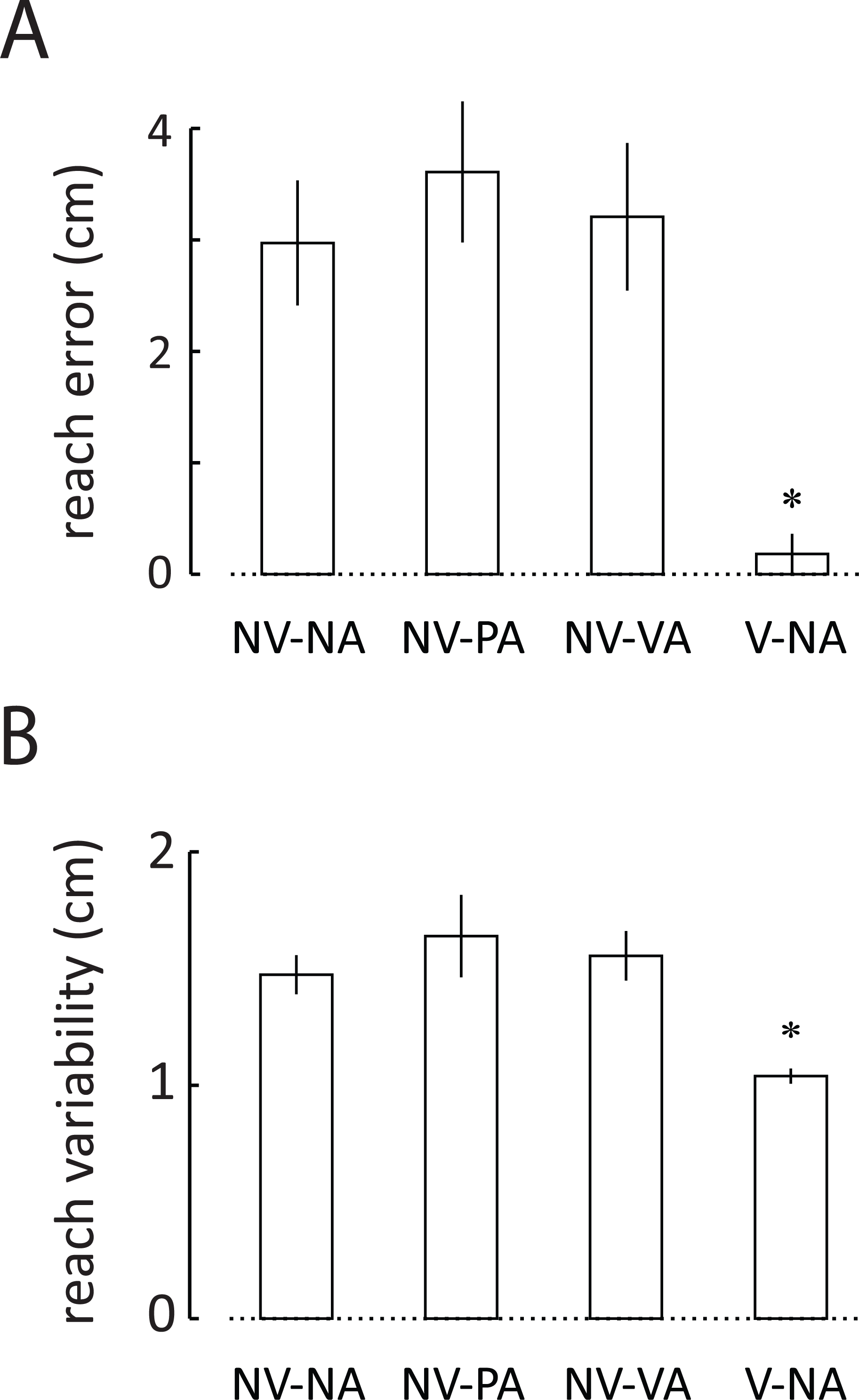
Average kinematic performance across the study cohort for all the measured conditions: A) mean reach error at peak movement extent. B) mean variability of reach at peak movement extent. Error bars: 1 standard error of the mean (SEM). Condition labels as for Figure 2. Testing conditions that differed systematically from the NV-NA control condition are indicated by an asterisk (p<0.05).

### Fidelity of self-assessment

To determine the extent to which memories recalled during the self-assessment tasks accurately reflected peak movement extents, we regressed the assessed movement extents 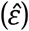 upon the actual movement extents (*ε*) for both the proprioceptive and visual self-assessment conditions (Figs 4A and 4B, respectively). In both cases, best-fit lines to the individual subject datasets reveal significant underestimation of assessed movement extents, as well as compression of the assessed range of extents relative to the actual range of movement extents. These individual results were characteristic of the cohort data. To assess the magnitude of bias between self-assessed and actual movement extents, we used two-sided, one-sample *t*-test to compare the y intercepts of the best fit lines from the individual subject data to a value of zero. In both self-assessment conditions, the subjects’ reported movement extents significantly underestimated the actual reach extents (PA: *t*_19_ = 2.65; p = 0.016; VA: *t*_19_ = 3.84; p = 0.001). Using two-sided, one-sample *t*-test to compare (to a value of 1) the slopes of the best fit lines from the individual subject data, we found that the slope was significantly less than unity for both self-assessment conditions (PA: *t*_19_ = 5.30; p < 0.0005; VA: *t*_19_ = 9.26; p < 0.0005). These results demonstrate that across our study cohort, declarative memories of the most recent movement extents differed systematically from the actual movement extents in both bias and range.

**Figure 4:**
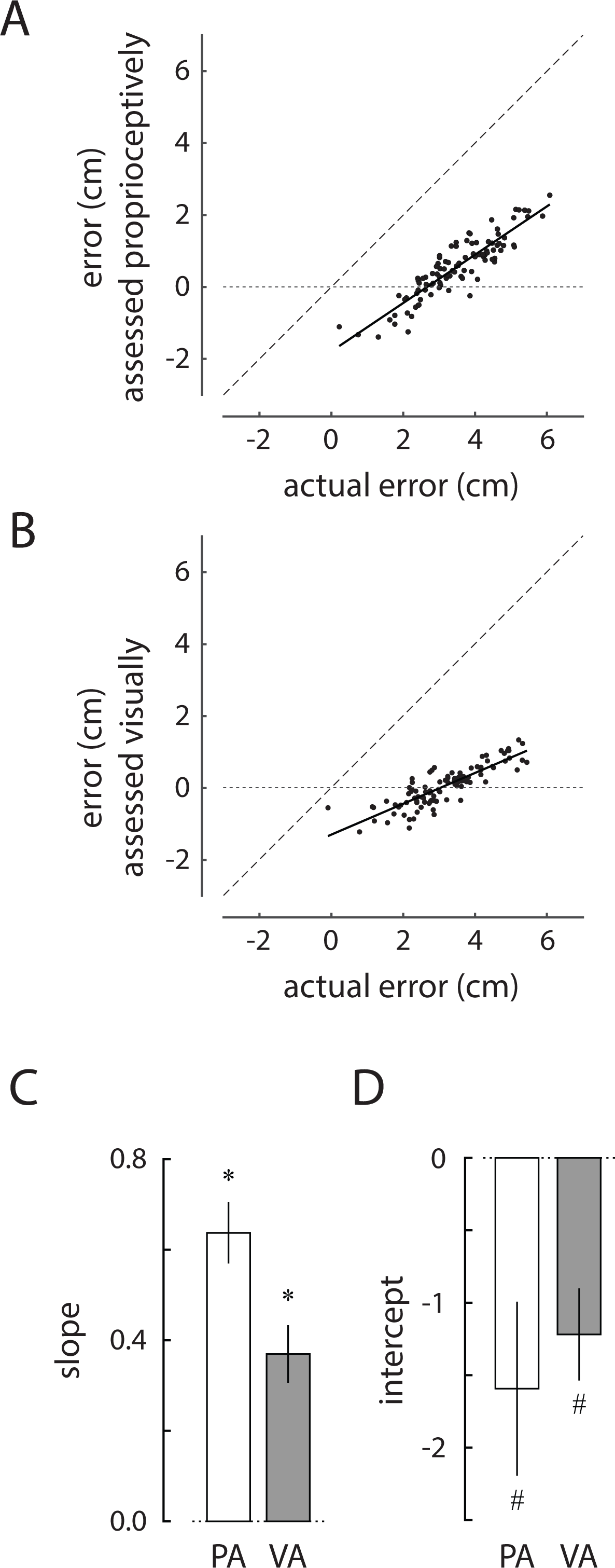
Comparison of the 100 post-reach, self-reported movement extents to the actual movement extents in the A) NV-PA and B) NV-VA trial blocks for a selected subject. The thick solid line is the line of best fit to the data. A dashed line with unity slope is shown as a guide. C) Cohort results: slope of the relationship between self-assessed and actual reach extents in the two self-assessment trial blocks: White bars: actual reach extents; Grey bars: self-assessed reach extents. Here, asterisks (*) indicate conditions wherein the population statistics differed significantly from unity slope (p<0.05). D) Cohort results for mean offset in the relationship between self-assessed and actual reach extents in the two self-assessment blocks. Here, hash signs (#) indicate conditions wherein the population statistics differed significantly from an offset of 0 (p<0.05).

To evaluate whether self-assessments of hand movement extent depended on the reporting mode (PA or VA), we compared the extent to which bias and slope values differed across the two self-assessment conditions. While paired *t*-test found no significant difference in bias between the two self-assessment blocks (*t*_19_ = 0.81; p < 0.430), the slope value in the PA block (0.64 ± 0.30) was significantly greater (and closer to the ideal value of 1.0) than slope values in the VA block (0.37 ± 0.29) (paired *t*-test: *t*_19_ = 4.27; p < 0.0005). Thus, the two modes of self-assessment differed from each other, with proprioceptive assessments (the within-modality reporting condition) yielding more accurate but nevertheless biased representations of actual movement extents than visual assessments (the cross-modality reporting condition).

### Contribution of implicit and explicit memories to sensorimotor adaptation of reach extent

To evaluate the extent to which implicit and explicit memories of recent reach performance contribute to sensorimotor adaptation of reach extent, we used a pair of stepwise linear regression analyses to fit a family of linear adaptation models to the series of reach errors (*ε*_*i*,_) within each self-assessment block of trials. These models included input terms that contain information about the original load sequence *k*_*i*_ and memory of the most recent load *k*_*i*,-1_. Also included were implicit and explicit memory terms (*ei*-1 and *ê*_*i*-1_ respectively); recall that by construction, these terms contain novel information about past kinematic performance that is unrelated to the trial-by-trial changes in robotic load. For the proprioceptive self-assessment block (NV-PA; Table 1), the model structure that most parsimoniously explained the data included only those terms related to the robotic load, a memory of the previous trial’s load, and an (implicit) memory of the previous trial’s movement extent (i.e., *εi*, ∼ *k*_*i*_ + *k*_*i*-1_ + *e*_*i*-1_). Whereas ∼57% of the trial-by-trial variations in reach extent were predicted by concurrent variations in the robot’s load, ∼33% and ∼4% of the variations in reach extent were predicted respectively by variations in the implicit memory of kinematic performance *e*_*i*-1_, and by prior robotic load *k*_*i*-1_. Adding the explicit memory term *ê*_*i*-1_ improved the model’s predictive ability (R2) by only 0.1%. This increment was neither significant nor meaningful. A similar outcome was obtained from the stepwise regression analysis applied to data from the visual self-assessment block (NV-VA; Table 2). In that case, ∼69% of the trial-by-trial variations in reach extent were predicted by variations in the robot’s load on the same trial, whereas ∼17% and 9% of the variations in reach extent were predicted by variations in the implicit memory of kinematic performance *e*_*i*-1_ and prior robotic load *k*_*i*-1_. Adding the explicit memory term *ê*_*i*-1_ improved the model’s predictive ability (R2) by only 0.2%. While this increment was marginally significant in the statistical sense (p = 0.045), the contribution of the explicit memory component *ê*_*i*-1_ to overall model performance was nearly 2 orders of magnitude smaller than that of the implicit memory component *e*_*i*-1_.

**Table 1:** Results of Stepwise Regression Analysis – NV-PA Condition

| Model Structure | $R^2$ | $\Delta R^2$ | $\Delta F$ | Sig. of $\Delta F$ |
| --- | --- | --- | --- | --- |
| $\varepsilon_i \sim k_i$ | 0.568 | 0.568 | 129.027 | 0.000 |
| $\varepsilon_i \sim k_i + e_{i-1}$ | 0.897 | 0.329 | 309.885 | 0.000 |
| * $\varepsilon_i \sim k_i + k_{i-1} + e_{i-1}$ | 0.934 | 0.037 | 53.769 | 0.000 |
| $\varepsilon_i \sim k_i + k_{i-1} + e_{i-1} + \hat{e}_{i-1}$ | 0.935 | 0.001 | 0.827 | 0.365 |

Row 3 (\*) corresponds to the most parsimonious model identified using stepwise regression analysis.
Note: The last row in this table shows the impact of adding a term that included information unique to explicit memory of performance. This term did not contribute significantly (i.e., $p > 0.05$ ) or meaningfully to the overall model performance $R^2$ .

**Table 2:** Results of Stepwise Regression Analysis – NV-VA Condition

| Model Structure | $R^2$ | $\Delta R^2$ | $\Delta F$ | Sig. of $\Delta F$ |
| --- | --- | --- | --- | --- |
| $\varepsilon_i \sim k_i$ | 0.690 | 0.690 | 218.105 | 0.000 |
| $\varepsilon_i \sim k_i + e_{i-1}$ | 0.855 | 0.165 | 110.685 | 0.000 |
| * $\varepsilon_i \sim k_i + k_{i-1} + e_{i-1}$ | 0.948 | 0.092 | 169.657 | 0.000 |
| $\varepsilon_i \sim k_i + k_{i-1} + e_{i-1} + \hat{e}_{i-1}$ | 0.950 | 0.002 | 4.116 | 0.045 |

Row 4 corresponds to the best model identified using stepwise regression analysis. However, adding a term that included information unique to explicit memory of performance yielded only marginally significant performance improvement (i.e., $p = 0.045$ ) such that this term did not contribute meaningfully to the overall model performance $R^2$ . Thus, Row 3 (\*) is considered the most parsimonious model.

We obtained similar results when we repeated the stepwise analyses using the ‘raw’ kinematic memories *ε*_*i*-1_and 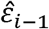 as inputs rather than the sequences *e*_*i*-1_ and *ê*_*i*-1_, which were adjusted to remove multicollinearity with *k*_*i*-1_ (details not presented). Finally, we obtained identical results in analyses that also corrected for collinearity between *ε*_*i*-1_and 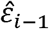, as was shown in Figures 4A and 4B. In all cases, the contribution of the explicit memory component to overall model performance was negligible compared to that of the implicit memory component. The parsimonious model structure, identified in the self-assessment blocks, is consistent with the results of prior modeling of memory-based motor adaptation during horizontal planar reaching (i.e., Eq 1; Scheidt et al., 2001; Takahashi et al., 2001; Judkins and Scheidt, 2014). In each case, the only terms that contribute meaningfully when adapting to rapidly changing mechanical environments are those related to the robotic load, memory of the previous trial’s load, and implicit memory of the previous trial’s movement extent.

To test this conclusion, we performed a head-to-head comparison of two models based on Eq 1 that differed only in that one model additionally included an explicit memory term 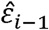 We used a cross-validation approach to compare the ability of the two models to capture the variability of the individual subject performances in the two self-assessment conditions. In each case, we separately fit the two models to one half of each participant’s dataset (the parameter identification dataset) and assessed each model’s ability to predict performance in the other half (the validation dataset). We used one-sided, paired *t*-test to determine whether the amount of data variance accounted for (VAF) would increase in the augmented model vs. that of Equation 1 due to the additional information about explicit memories provided by visual and/or proprioceptive self-assessment. In both self-assessment conditions, addition of an explicit memory term did not improve model performance (PA: *t*_19_ = -1.10; p = 0.857; VA: *t*_19_ = 0.833; p = 0.208) (Fig 5). Taken together, these results indicate that explicit memory systems do not contribute significantly to sensorimotor adaptation of movement extent during goal-directed reaching.

**Figure 5:**
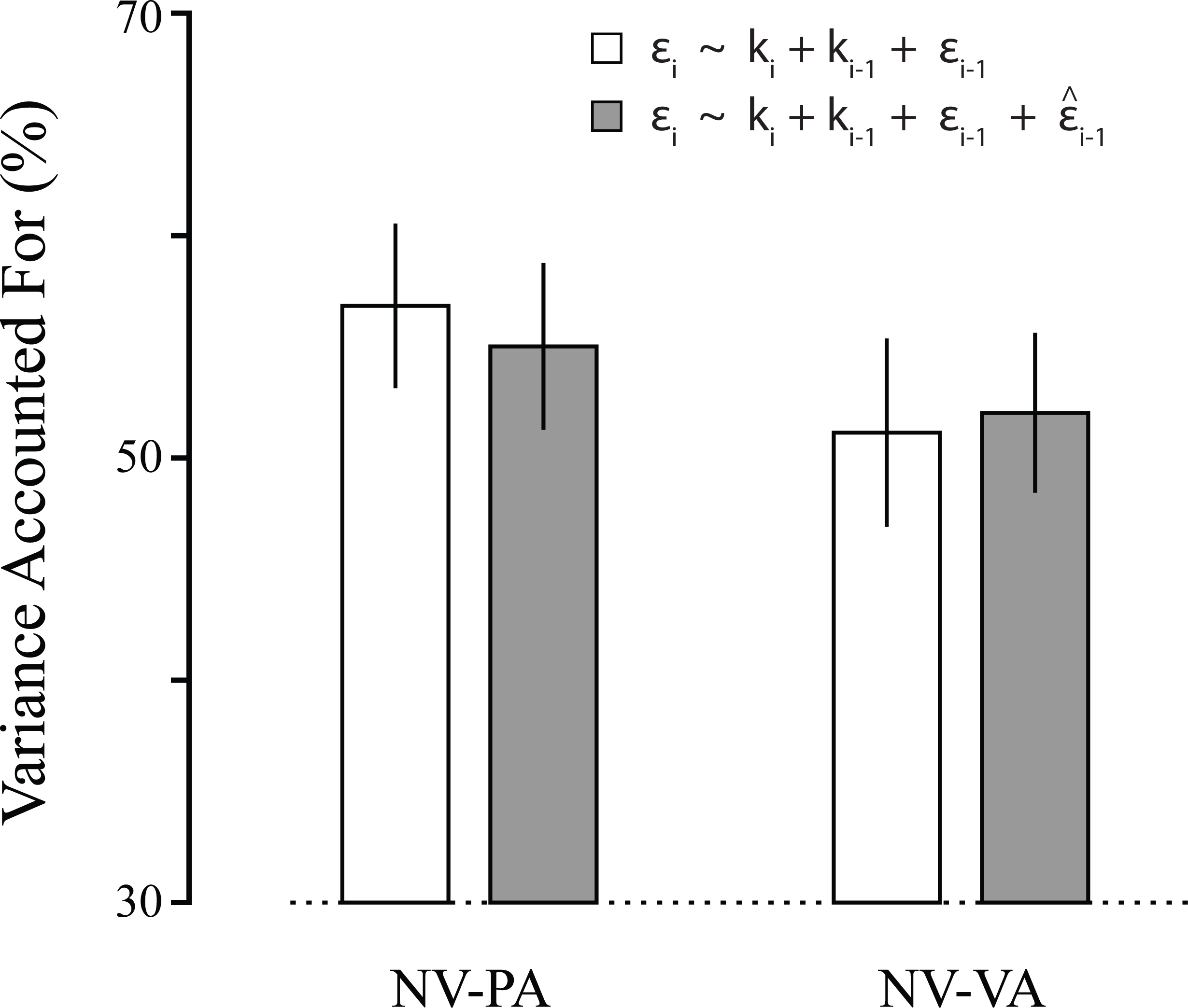
Results of the cross-validation analysis. Data variance accounted for (VAF) by the two adaptation models described in the main text. The two models were separately fit to one half of each participant’s dataset in both the NV-PA and NV-VA trial blocks. They models were then assessed for their ability to predict the data in the other half of the datasets. White bars: the model of Equation 1, which includes only implicit sensorimotor memories; Grey bars: the augmented model, which additionally includes the explicit (declarative, self-assessed) sensorimotor memories.

### Impact of vision and self-assessment on sensorimotor adaptation - model coefficients

We examined how the different testing conditions might impact the extent to which implicit memories contribute to sensorimotor adaptation to changing environmental loads. Justified by the stepwise regression analyses, we fit the model of Eq 1 to each subject’s time series of reach error (*ε*_*i*_) and spring stiffness (*k*_*i*_) values within each of the four trial blocks. We compared the resulting model coefficients (*b*_0_, *b*_1_, and *a*_1_) within the test and contrast conditions to those in the control condition to determine the relative impact of interposed self-assessments on the identified sensorimotor information filters (Figure 6A-C). One-way repeated measures ANOVA found a main effect of testing condition on each of the model parameters [F_(3,57)_ ≥ 6.04, p ≤ 0.001 in each case]. Similar to what was reported for reach kinematics in Figure 3, the observed effects were due to the presence of visual feedback of cursor motion, not to the interposed self-assessments. Post hoc *t*-tests confirmed that *b*_0_, *b*_1_, and *a*_1_ values in the V-NA contrast condition were smaller than those in the NV-NA control condition (*t*_19_ ≥ 2.94; p ≤ 0.009 in each case). By contrast, we observed no systematic impact of interposed self-assessment on these parameters. The *b*_0_, *b*_1_, and *a*_1_ values in the two self-assessment blocks did not differ systematically from those in the NV-NA control condition (*t*_19_ ≤ 1.33; p ≥ 0.20 in all cases). Of particular interest for this study is model parameter *a*1, which determines how implicit memories of prior reach errors influence subsequent reach performance. While providing a visual representation of the moving hand significantly impacted how sensorimotor memories are used to adapt reaches from one trial to the next, interposing self-assessments between reaches did not have significant impact.

**Figure 6:**
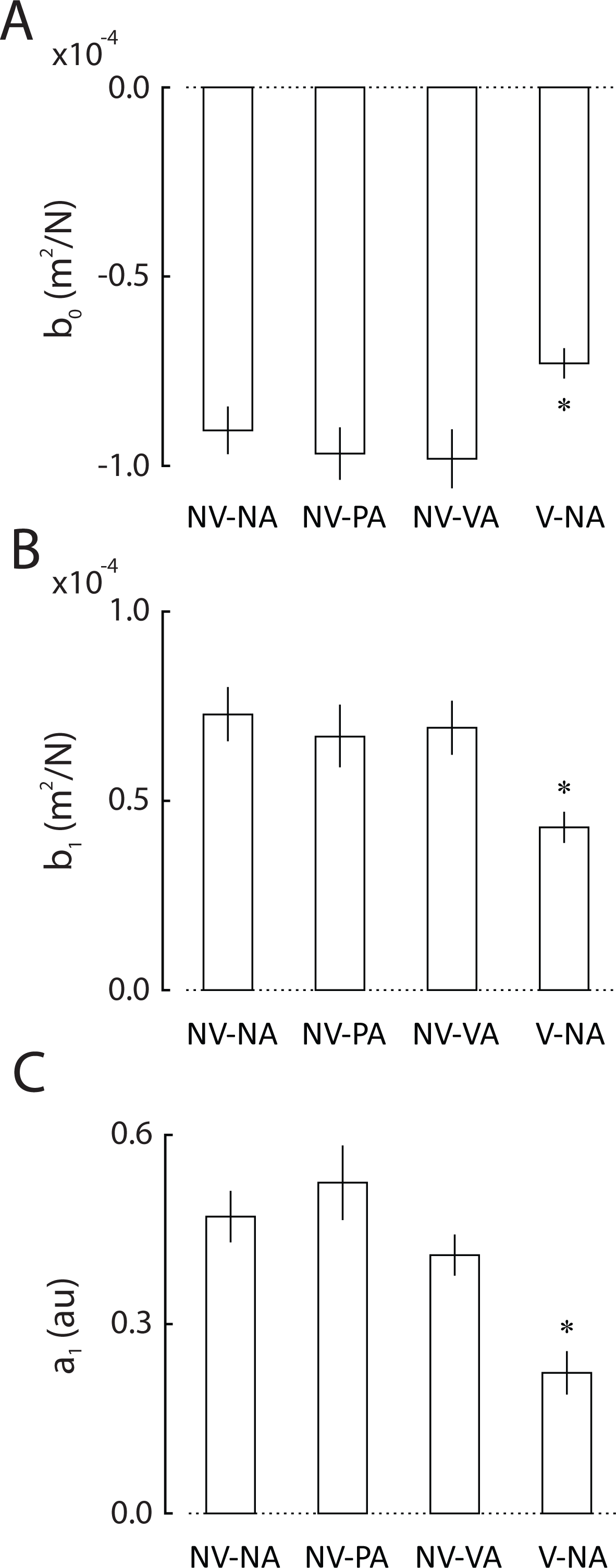
Adaptation model coefficients (Eq 1) across all four testing conditions for the study cohort. A) Model coefficient *b*_*0*_ vs. testing condition. Condition labels as for Figure 2. Error bars: ±1 SEM. B) Model coefficient *b*_1_ vs. testing condition. C) Model coefficient *a*_1_ vs. testing condition. Testing conditions that differed systematically from the NV-NA control condition are indicated by an asterisk (p<0.05).

## Discussion

We tested how implicit and explicit memories of recent kinematic performance contribute to sensorimotor adaptation of movement extent during goal-directed reaching. To gain insight into the subject’s explicit memories of kinematic performance, we sometimes required subjects to recall and report the hand’s location at the moment of peak movement extent. These self-assessments were performed visual or kinematically. We assessed the effects of these self-assessments on a control (non-visual feedback) condition, and compared the findings to those obtained in a contrast condition where visual feedback of cursor motion was provided. We then assessed the effects of interposed self-assessment on trial-by-trial motor adaptation under the different reporting conditions.

Within the study cohort, movement kinematics were similar across the testing conditions (Fig 2). Relative to the no-assessment control condition, interposing self-assessments between successive reaches neither improved nor degraded reach kinematics (Fig 3), nor did it alter the relative contributions of implicit memories of prior performance features to sensorimotor adaptation (Fig 6). By contrast, and consistent with prior research (Judkins and Scheidt 2014), providing concurrent visual feedback of cursor motion allowed subjects to move more accurately and precisely (Fig 3) and they did so by using sensorimotor memories differently relative to the control condition (Figs 6). While our test of how people use memories is sensitive to differences in sensory feedback during movement, we found no discernible effect of interposed self-assessments. Thus, interposing self-assessments between successive reaches provided access to explicit memories of reach performance without significantly impacting how implicit memories are used to adapt to changing environmental loads.

We next tested whether implicit and explicit memories draw upon a common representation of kinematic performance during repetitive goal-directed reaching. We considered actual and recalled errors as proxies for implicit and explicit memories, respectively. Consistent with prior literature supporting the idea that implicit and explicit memory systems are distinct (Corkin 1984; Keisler and Shadmehr 2010; Eichenbaum 2013; Taylor et al. 2014; McDougle et al. 2015) we found marked differences between actual and reported movement extents in terms of the offset (bias) and range (slope) of the linear relation between these two variables (Fig 4). While this was true for both the visual and proprioceptive self-assessment testing blocks, explicit recall and reporting was better using manual (proprioceptive) reporting vs. visual reporting in the sense that the slope of the relationship between actual and reported movement extents was closer to the ideal value of 1.0 using manual reporting. These results support the conclusion that implicit and explicit recall of movement extent draw upon distinct memories of reach performance.

Finally, we evaluated the extent to which implicit and explicit memories of recent reach performance contribute to sensorimotor adaptation of movement extent. Stepwise regression analysis found that adding an explicit memory term added negligibly to the ability of a simpler model (Eq 1) to predict reach performance when adapting to rapidly changing mechanical environments; the contribution of the explicit memory component to overall model performance was nearly 2 orders of magnitude smaller than that of the implicit memory component. We also used a cross-validation approach to compare the ability of models with and without an explicit memory term to capture the variability of the individual subject performances in the two self-assessment conditions. Contrary to the expectation that a model including both explicit and implicit memory terms would outperform a model that includes only implicit memories, the model with the explicit memory term fared no better than the simpler model of Eq 1 (Fig 5). Taken together, our results indicate that sensorimotor adaptation of reach extent recruits predominantly implicit sensorimotor memories; explicit memories do not contribute meaningfully.

### Separate implicit and explicit memory systems

Studies of amnesic patients, including patient H.M., indicate that there exist distinct memory systems serving short-term and procedural memories necessary for implicit perceptual and motor skill acquisition (Corkin 1984), and those serving declarative/episodic memories required for the explicit and conscious recollection of prior events (Eichenbaum 2013). When neurologically intact subjects are exposed to a novel visuomotor perturbation during reaching, implicit and explicit processes can both contribute to the adaptive response (Mazzoni and Krakauer 2006; Taylor et al. 2014; McDougle et al. 2015). While these systems are distinct (c.f., Redding et al. 2005), they can interact in unexpected ways. In one study, Mazzoni and Krakauer (2006) asked 2 groups of subjects to use wrist movements to move a cursor to visual targets spaced 45° apart around a central starting position in the 2D plane of a computer screen. On some trials, a visuomotor rotation was experimentally imposed such that the cursor moved 45° counterclockwise (CCW) about the center of the starting location. One group was instructed to strategically counter-rotate their wrist motions to compensate for the imposed rotation by aiming for the neighboring clockwise target. Whereas instructed strategic re-aiming was initially effective in cancelling the imposed rotation, subjects unexpectedly made increasingly large directional errors as they continued to make movements. Later, when these subjects were informed that the visuomotor rotation would be switched off, they nevertheless made persistent aftereffects demonstrating that implicit learning of the rotation had occurred despite application of the explicit counter-rotation strategy. The authors concluded that implicit adaptation continues even when participants are provided an effective explicit strategy to perform a movement.

In another study, Taylor and colleagues assayed the contributions of explicit strategic re-aiming and implicit sensorimotor adaptation to an imposed visuomotor rotation by asking subjects to verbally report the direction of their intended aim prior to each reach (Taylor et al. 2014). Subjects made center-out reaches to each of 8 visually displayed targets in the horizontal plane using a digitizing tablet and pen. After first making several center-out reaches to each target, a 45° CCW visuomotor rotation was imposed on the motion of a cursor. During these rotation trials, half of the subjects were asked to report their intended aim prior to each reach using a ring of numbered landmarks displayed just above the plane of hand motion. To calculate the magnitude of implicit adaptation on any given trial, the investigators subtracted the intended aim (verbally reported) from the observed heading angle of the hand. The remaining subjects served as controls and were not required to report the intended aim. Target capture errors asymptotically approached zero well within the block of 300 trials. Although the angle of aim averaged ∼30° early in the rotation block, it fell to ∼15° by the end of the block. Accordingly, the remaining ∼30° of compensation was assumed to be due to implicit learning. Taken together, the results provide compelling evidence for both implicit and explicit contributions to learning in a visuomotor adaptation task. Explicit re-aiming played a more dominant role early in exposure and implicit learning played a more dominant role later. Mixed contributions of implicit and explicit processes have also been reported in studies of force field adaptation (Keisler and Shadmehr 2010).

Considering these prior works, the results of our study are surprising. Whereas people evidently re-aim subsequent reaching movements to compensate for directional errors recalled from previous movement attempts in visuomotor rotation studies, we found here that explicit memories contribute negligibly to the adaptive compensation for reach extent errors caused by unpredictable forces opposing movement. At least two main differences between the studies may have motivated the different findings.

The first difference relates to the type of movement perturbations subjects had to overcome and to the type of performance feedback provided during the experiments. In the studies by Taylor and colleagues (Taylor et al. 2014; McDougle et al. 2015), subjects were required to compensate for an imposed visuomotor rotation and were provided visual feedback of the hand’s heading direction during and/or after movement. No special efforts were made to eliminate or otherwise confound somatosensory sensation. By contrast, subjects in our study were required to compensate for environmental forces that impacted movement extent and the movements themselves were physically constrained to lie along the straight line connecting the starting position and the goal, thus eliminating direction errors. Feedback of movement extent was limited to intrinsic proprioceptive sensations in the relevant self-assessment blocks of our study. It is possible that the different outcomes reflect fundamental differences in the way the brain plans and controls movement direction and movement extent. Careful studies of movement accuracy, variability, and reaction time support the hypothesis that the direction and extent of reaching and pointing movements are specified via separate cognitive channels (Bock and Eckmiller 1986; Bock 1992; Gordon et al. 1994a; Ghez et al. 1997; Bhat and Sanes 1998; Vindras and Viviani 1998; Krakauer et al. 2000; Sainburg et al. 2003). While these channels might operate in parallel to some degree, requiring subjects to re-aim a movement appears to delay and prolong extent specification in a way that depends on whether or not the desired movement is predictable or unpredictable and on the amount of time allowed for planning (Ghez et al. 1997; Bhat and Sanes 1998). Processes that compensate for a visuomotor rotation may engage mental computations that counter-rotate the intended hand movement through a series of intermediate movement directions (cf., Georgopoulos and Massey 1987). By contrast, learning of a movement gain (Bock 1992; Pine et al. 1996) may involve the learning of a global scaling factor (i.e., *vigor*; c.f., Summerside et al. 2018) that makes modest demands on short-term working memory (Pine et al. 1996). We suggest that the mental rotations needed to compensate a visuomotor rotation may engage an explicit strategy of re-aiming, at least during initial exposure to the perturbation, whereas adapting movement extent may engage short-term working memory in a way that is not readily accessible to conscious recall and verbal description.

The second difference pertains to the relative predictability of the perturbations imposed in the different studies. Whereas the earlier cited studies invoke a predictable visuomotor rotation (Mazzoni and Krakauer 2006; Taylor et al. 2014; McDougle et al. 2015), here we required subjects to adapt to an unpredictable series of spring-like loads opposing motion. It seems reasonable that an explicit re-aiming strategy would accrue and be refined with repeated exposure to the same perturbation as in the visuomotor rotation studies. We suggest that strategic compensation was discouraged in our study because the unpredictable perturbations induced large and persistently variable errors. It seems highly unlikely that subjects would consciously choose - consistently and without guidance - a strategy of multimodal sensory integration that draws upon memories of prior hand forces and movement extent errors in the specific combination described by Eq 1. This outcome was insensitive to whether subjects reported their recalled performance visually or proprioceptively, or not at all. When asked after completing the experiments, no subject in our study was able to verbalize any consistent strategy used to acquire the target from one trial to the next. Thus, our experimental approach engaged implicit adaptation to the practical exclusion of explicit strategic compensation.

### Intent matters: The functional independence of reaching and pointing

The proprioceptive self-assessment block required subjects to move their hand out-and-back to capture a target at the peak of the movement, and then to point at the remembered reversal location with the same, moving hand. Many previous studies have reported that sensorimotor adaptation of goal directed movements is a limited memory process wherein errors on the most recent movements influence plans for subsequent movements such that errors are reduced (Thoroughman and Shadmehr 2000; Scheidt et al. 2001; Smith et al. 2006; Lee and Schweighofer 2009). Interference between interleaved point-to-point and out-and-back target capture movements directed to the same target has previously been reported (Scheidt and Ghez 2007). In that study, out-and-back target capture movements overshot the target when performed immediately after performing a point-to-point reach to the same spatial target. The authors argued that such errors were the result of violated expectations, in that stabilizing muscle co-contractions at the end of accurate point-to-point movements caused elevated joint viscoelasticity that was not present near the target during the out-and-back movements. Engaging the same plan to initiate both kinds of movements led to the observed overshoot of out-and-back movements performed after point-to-point reaches (Scheidt and Ghez 2007; Scheidt et al. 2011). One might reasonably expect therefore to observe interference between pointing movements and out-and-back target capture movements in the current study because proprioceptive self-assessment required hand movement to the same spatial location as the preceding out-and-back reach. Nevertheless, interposed pointing had no significant impact on movement kinematics or on how participants used implicit memories to adjust subsequent out-and-back movements to compensate for changing loads in the current study. Kinematic performance and adaptation modeling in the NV-VA and NV-PA blocks were similar in all respects to those in the NV-NA block. By contrast, providing cursor feedback of ongoing movement changed not only kinematic performance, but also how memories were used from one trial to the next to compensate for unpredictable environmental loads. These last findings largely replicate those of Judkins and Scheidt (2014), who showed that providing ongoing visual feedback of endpoint movement improves the ability to reduce performance fluctuations caused by environmental loads that change rapidly from trial to trial.

Our inability to find impact of self-assessment was not the result of insensitivity of our experimental approach to quantifying how memories contribute to sensorimotor adaptation. Rather, our results likely reflect important differences between the reaching and pointing tasks employed in the current study. The absence of an interaction between reaching and pointing was a general phenomenon because performances in both self-assessment blocks (NV-PA and NV-VA) did not differ from those in the NV-NA control condition, whereas performance in all three of those conditions differed markedly from performance in the V-NA contrast condition. We conclude therefore that reach attempts in the two self-assessment blocks drew exclusively upon memories from prior out-and-back reaches rather than on the prior pointing movements. One possible reason for this result is that there was no violation of expectations during pointing. Whether subjects indicated peak movement extent by moving a visual indicator via button presses or by moving that same hand, there was no performance error during pointing – and no sensory prediction error – because the indicating endpoint ended up exactly where the subject decided it should go. In the absence of error, no updating of the motor plan for the subsequent movement is expected.

### Limitations and future directions

Our study had at least two limitations. First and foremost, we narrowly focused on the contributions of explicit memory to the trial-by-trial adaptation of reach extent and did not broadly assess the extent to which the subject’s explicit plan for the upcoming movement’s vigor might have been consciously available. Taking inspiration from the studies of Taylor and colleagues (Taylor et al. 2014; McDougle et al. 2015), a future study could require subjects not only to recall and report their movement errors on the prior trial as done in the current study, but also to report how much they plan to change the vigor with which they expect to make the next movement. This could be reported on a subjective scale where a value of 0 indicates no change, whereas increases and decreases would range from very slight (±1) to the maximum possible (±10). By doing so, it would be possible to determine whether requiring subjects to articulate an explicit plan would change (i.e., interfere with) the nominal way in which subjects use implicit memories to update subsequent movements. Another limitation stems from the fact that we focused exclusively on kinematic memories. Equation 1 also includes a memory term related to the strength of the most recent robotic resistance to movement. A future study could independently confirm the results of the current study by asking subjects to recall and report the peak force experienced at the hand (i.e., *k*_*i*-1_) by having subjects manually replicate that peak force under isometric conditions in-between reach trials.

In conclusion, the experimental approach described here engaged implicit memories to compensate for reach extent errors, to the practical exclusion of explicit strategic compensation. Because interposed self-assessments provided access to explicit memories of reach performance without significantly impacting how implicit memories are used to adapt to changing environmental loads, a future neuroimaging study may be able to disentangle and visualize the different memory systems contributing to the conscious and subconscious responses to dynamically changing physical environments.

## Acknowledgements

This work was supported in part by a grant from the Marquette University Athletic and Human Performance Research Center, and by the National Science Foundation (NSF) under an Individual Research and Development Plan. Any opinions, findings, conclusions, or recommendations expressed in this material are those of the authors and do not necessarily reflect the views of the NSF.

